# Nociceptin/Orphanin FQ receptor agonism attenuates behavioral and neural responses to conditioned aversive stimuli

**DOI:** 10.1101/2025.06.05.658049

**Authors:** Kwang-Hyun Hur, Diego A. Pizzagalli, Jessi Stover, Kenroy Cayetano, Stephen J. Kohut

## Abstract

The nociceptin/orphanin FQ peptide (NOP) receptor has emerged as a promising anxiolytic target, as its activation has been shown to reduce anxiety-related behaviors in rodents. However, the mechanisms underlying these effects are not well understood. Here, we investigated the effects of the selective NOP receptor agonist SCH-221510 (SCH; 0.01-0.1 mg/kg, IM) on behavioral and neural responses to aversive stimuli in squirrel monkeys (n=3). Subjects underwent Pavlovian fear conditioning, wherein a visual conditioned stimulus (CS) was paired with the presentation of an aversive stimulus. Event-related fMRI was conducted in awake subjects to evaluate CS-evoked neural responses. Behavioral and neural responses to the CS were assessed across three experimental phases: pre-conditioning (Pre-C), post-conditioning (Post-C), and Post-C with SCH administration. In behavioral assessments, CS presentation during Post-C elicited a robust suppression of ongoing operant responding, which was absent during Pre-C and significantly attenuated by SCH treatment (0.1 mg/kg). Functional magnetic resonance imaging (fMRI) results revealed that, relative to Pre-C, CS presentation during Post-C was associated with increased BOLD activity in brain regions previously implicated in fear processing (e.g., amygdala), expression and regulation (e.g., prefrontal cortex; PFC), as well as sensory integration. Critically, SCH (0.1 mg/kg) administration significantly attenuated CS-induced neural activation in these regions. Furthermore, resting-state functional connectivity analysis revealed that SCH administration decreased connectivity between the PFC and the amygdala, while enhancing connectivity among subregions of the PFC. Collectively, these findings suggest that NOP receptor agonism may attenuate conditioned responses to aversive stimuli by modulating functional interactions within the PFC–amygdala circuit.

## 1. Introduction

Anxiety disorders are among the most prevalent psychiatric conditions, affecting approximately 300 million individuals worldwide [1]. Dysregulated or exaggerated fear responses have been recognized as significant risk factors for the development and maintenance of these disorders [2]. The neurobiological mechanisms underlying heightened fear-related anxiety have generally been characterized by maladaptive functional dynamics within prefrontal-limbic circuitry, including prefrontal cortex (PFC), amygdala, and hypothalamus, which have roles in fear processing and regulation [3, 4]. While the available pharmacological and psychotherapeutic interventions targeting these pathophysiological mechanisms provide symptomatic relief for some individuals, a substantial proportion of patients continue to experience treatment resistance or incomplete symptom remission [5]. These limitations underscore the critical need to identify alternative neuromodulatory systems involved in fear regulation, which may inform the development of more effective and mechanistically targeted interventions for anxiety disorders.

The nociceptin/orphanin FQ (N/OFQ) peptide (NOP) and its receptor system have emerged as a promising therapeutic target for anxiety disorders [6], given their high expression in fronto-limbic circuits [7]. Preclinical studies have demonstrated that activation of the NOP receptor effectively attenuates anxiety-related behaviors, including fear-potentiated startle, in rodent models [8, 9]. Despite these promising findings, the precise neurobiological mechanisms through which NOP modulates fear and anxiety remain incompletely understood. Specifically, the influence of NOP system activation on fear-related neural circuits, including the activity of key brain regions and the functional connectivity within these circuits, has not been characterized and warrants further investigation. Addressing these gaps in knowledge may provide new insights into the neurobiological underpinnings of fear-related anxiety disorders and facilitate the development of innovative therapeutic strategies targeting the NOP system.

Event-related functional magnetic resonance imaging (fMRI) has been widely employed in clinical research to investigate the neural correlates of anxiety, particularly in relation to maladaptive fear responses [10, 11]. This technique allows for high temporal resolution in capturing brain activity in response to discrete emotional or conditioned stimuli, making it especially suitable for probing the dynamic engagement of neural circuits involved in fear processing and regulation. Importantly, event-related fMRI has been used as a translational tool to evaluate the efficacy of various anxiolytic compounds by quantifying their modulatory effects on neural responses to aversive or threat-related stimuli [12]. Accordingly, this approach provides a rigorous framework for elucidating the neurobiological mechanisms by which candidate therapeutics, such as NOP receptor agonism, modulate fear-related circuit dynamics to exert anxiolytic effects.

In this study, we investigated the role of the NOP system in modulating fear-related behaviors and associated neural signatures using nonhuman primates (NHPs), given the substantial neuroanatomical homology between NHPs and humans [13], including the CNS distribution of NOP receptors [7, 14]. A Pavlovian fear conditioning procedure, integrated with a food-maintained operant responding task, was implemented to develop behavioral and neural responses to fear-related aversive stimuli [15]. This approach provides a translationally relevant measure of anxiety-related behavioral inhibition, capturing the extent to which conditioned aversive stimuli disrupt goal-directed behavior, which is frequently impaired in individuals with anxiety disorders [16]. Event-related fMRI was employed to investigate the neuronal mechanisms underlying NOP-mediated modulation of fear responses. By integrating behavioral and neuroimaging approaches, this study aims to delineate the circuit-level mechanisms by which NOP signaling regulates fear responses to aversive stimuli. Extrapolating from rodent findings, we hypothesized that NOP receptor agonism would attenuate behavioral and fMRI indices of fear responses.

## 2. Materials and Methods

### 2.1. Subjects

One male and two female adult squirrel monkeys (Saimiri sciureus) were used. A power analysis, grounded in prior studies from both our laboratory and other relevant research, confirmed that a sample size of three subjects was sufficient to achieve the statistical power necessary to detect significant effects in both behavioral and neuroimaging components of the study [17, 18]. Each subject served as its own control in a within-subject, repeated measures experimental design, ensuring reliable group-level effects [19]. Subjects were housed in stainless-steel cages within a temperature-and humidity-controlled vivarium, with a 12-hour light/dark cycle (lights on from 07:00 to 19:00). Monkeys were maintained at approximate free-feeding weights through daily feedings of a nutritionally balanced diet comprised of high-protein chow (#5040, LabDiet, St. Louis, MO, USA) along with fresh fruits and vegetables. Water was available *ad libitum* outside of the experimental session. An assortment of manipulable toys, mirrors, puzzles, and background music during light hours were provided as part of an environmental enrichment program. Animal care and research were conducted according to the guidelines provided by the Institute of Laboratory Animal Resources and the National Institutes of Health Office of Laboratory Animal Welfare. The facility is licensed by the U.S. Department of Agriculture, and all experimental protocols were approved by the Institutional Animal Care and Use Committee at McLean Hospital.

### 2.2. Drugs

The nociceptin/orphanin FQ peptide (NOP) receptor agonist SCH-221510 (SCH, Tocris, Ellisville, MO, USA) was prepared in a vehicle solution consisting of 10% DMSO, 10% Tween 80, and 80% saline. SCH or vehicle was administered via intramuscular (IM) injection into the thigh muscle at a volume not exceeding 0.3 mL/kg of body weight, 30 min prior to behavioral assessments or MRI scans. The dose range and pretreatment interval were determined on the basis of previous studies [20, 21].

### 2.3. Behavioral Procedures

A multi-phase procedure was implemented, consisting of operant behavioral training to establish task performance, which was followed by Pavlovian fear conditioning. This protocol integrated Skinnerian operant conditioning principles within a fear-related paradigm [15, 22], enabling the assessment of conditioned suppression of operant behavior in response to presentation of visual stimuli associated with aversive stimuli. All experimental procedures, including the presentation of stimuli, detection of operant responses, delivery of milk reinforcement, application of electrical stimuli, and data acquisition, were controlled and recorded using Med Associates software (MedPC v4.2; Med Associates, St. Albans, VT, USA).

#### 2.3.1. Operant Behavioral Training

Subjects were positioned in a prone posture within a custom-designed MR-compatible chair, located inside a sound-attenuated and ventilated chamber [23]. A light panel with three distinct light-emitting diodes (LED) (green: left, yellow: center, red: right) was placed 20 cm in front of the subject’s face. A customized lickometer, which also functioned as a milk delivery tube, was positioned within tongue-reach proximity to facilitate operant responses. Each session commenced with the activation of the house light, signaling the start of a 300-sec session. The center light on the panel was illuminated to indicate the availability of an operant response opportunity. The center light remained active for 30-sec, during which subjects were required to perform a fixed ratio (FR) of operant responses (licks) on the lickometer, with the FR requirement varying across individuals (1-5). Completion of the required FR response resulted in the center light turning off and the delivery of 0.3 mL of 30% condensed milk as a reinforcer, followed by a 25-sec short timeout period. If the subject failed to meet the FR response requirement within the 30-sec window, the center light was turned off, and a 25-sec short-timeout period ensued without milk delivery. The conclusion of each 300-sec session was marked by the deactivation of the house light. Following a 90-sec long-timeout period, the house light reactivated to signal the start of a new session, with a total of four sessions conducted per day. Operant behavioral training continued until subjects achieved a consistent rate of ≥ 90% in completing the FR requirement for three consecutive days.

#### 2.3.2. Pavlovian Fear Conditioning

Upon completion of operant behavioral training, subjects underwent Pavlovian fear conditioning to associate a specific visual cue (conditioned stimulus, CS) with an aversive unconditioned stimulus (US; electric stimulation). During the 300-sec session, subjects were allowed to engage in operant responding (i.e., licking the lickometer) to obtain a milk reinforcement, consistent with the operant behavioral training phase. The CS, operationalized as the illumination of either the left or right panel LED (counterbalanced across subjects), was presented for 60-sec at a randomized timepoint within the session to mitigate temporal predictability, except during test sessions, where it was presented within a predetermined 120–180-sec time window. During CS^+^ trials, an electrical pulse (US; 0.5 mA, 0.2 sec) was delivered to the subject’s tail at a random time within the 60-sec CS^+^ period to establish an aversive association. Conversely, during CS^-^ trials (illumination of the remaining panel light), no stimulation was delivered, serving as a control condition. Each conditioning day consisted of four sessions: two baseline sessions (no CS presentations), one CS^+^ session, and one CS^-^ session. The sequence of sessions was counterbalanced across days and subjects to mitigate potential order effects. Fear conditioning was assessed by measuring conditioned suppression of operant responses during CS^+^ trials, operationally defined as a reduction in response rate to less than 25% of the baseline level. The 120-sec pre-CS period served as the baseline to avoid potential confounding effects of the CS presentation. Notably, in CS^+^ sessions following fear conditioning, responses during the post-CS period were lower than during the pre-CS period (Figure S1A). Response rate was calculated using the following equation: (number of responses during the CS period / 60-sec) / (number of responses during the pre-CS period / 120-sec) × 100 and is expressed as a percentage (%). After the fear conditioning phase, subjects underwent test sessions to evaluate the effects of NOP agonist (SCH 0.01-0.1mg/kg) pretreatment on conditioned behavioral responses to CS presentation. The effective drug dose for MRI procedures was determined based on observed significant behavioral effects. Each drug test was conducted with at least a three-day interval, and suppression of operant responses to the CS^+^ was confirmed prior to each test.

### 2.4. Magnetic Resonance Imaging (MRI)

A cue-reactivity paradigm incorporating event-related fMRI in awake subjects, based on those procedures used in clinical studies [10, 11], was utilized to investigate neural responses to conditioned aversive cues, with a particular focus on their association with the NOP system. During each fMRI session, electrical stimulation-associated (CS^+^) and control (CS^-^) light stimuli were presented in a pseudorandomized sequence to prevent anticipatory responses and ensure sustained task engagement. No electrical stimulations were delivered during the fMRI session, allowing for the isolated assessment of neural responses elicited by the conditioned cues. Each CS was presented for 6-sec, with an interstimulus interval jittered between 15-and 30-sec, totaling 23 presentations for each CS. The light presentation time and interval were chosen based on previous clinical cue-reactivity studies [24], and considering that the hemodynamic response to brief neuronal activity usually reaches its peak within 5–8-sec and returns to baseline within 15–30-sec [25, 26]. Each subject underwent MRI scans across three experimental phases: pre-conditioning (Pre-C), post-conditioning (Post-C), and post-conditioning with NOP agonist (SCH, 0.1 mg/kg) administration (Post-C+SCH). During the Pre-C phase, baseline neural activity in response to each CS was assessed to establish a reference for subsequent analyses. In the Post-C phase, conditioning-induced alterations in neural activation were examined by comparing responses to each CS before and after Pavlovian fear conditioning. Finally, in the Post-C+SCH phase, the modulatory effects of NOP signaling on fear-related neural dynamics were evaluated by examining drug-induced changes in neural responses to the CSs. Prior to the Post-C and Post-C+SCH phase scans, behavioral suppression under CS^+^ presentation was confirmed.

#### 2.4.1. Acclimation and Training for Awake MRI

To ensure subjects’ compliance and comfort during awake MRI scanning, an extensive behavioral training protocol was implemented, following a gradual adaptation protocol previously established in our laboratory [23]. During acclimation sessions, shaping techniques with milk reinforcement were employed to train subjects to remain in a prone position within a custom-designed, MR-compatible chair for brief periods of 5-10 mins. The session duration was progressively increased to 30 mins over successive training sessions. Following successful habituation to the chair, subjects were introduced to a 3D-printed head-positioning helmet, specifically designed using high-resolution anatomical images of squirrel monkeys. The helmet was padded to enhance comfort while effectively minimizing head motion and was equipped with a mount for a transmit/receive surface coil. The helmet was securely affixed to the chair using nylon screws to ensure stability. Once subjects were habituated to the helmet, the session duration was further extended to 60 mins, equivalent to the expected duration of the MRI session. In the final phase of acclimation, subjects were exposed to an MRI simulator housed within the laboratory. Recorded sounds of echo-planar imaging (EPI) sequences, played at decibel levels (∼90–100 dB) comparable to those experienced within the actual MRI bore, were introduced to simulate the scanning environment. Subjects were continuously monitored throughout the training sessions via live video feeds (VID-CAM-MONO-1 with SOF-842; Med-Associates, St. Albans, VT, USA). Vital signs, including heart rate and oxygen saturation (SpO₂), were measured using the Nonin 2500 pulse oximeter (Nonin Medical Inc., Plymouth, MN, USA). To maintain procedural familiarity, subjects underwent weekly training sessions throughout the study.

#### 2.4.2. MRI Data Acquisition and Preprocessing

All MRI data were acquired using a 9.4 Tesla horizontal bore magnet system (400 mm diameter; Varian Inc., Palo Alto, CA, USA) at McLean Hospital. The system was equipped with an 11.6 cm inner-diameter gradient coil capable of producing a maximum gradient strength of 45 G/cm. A single-loop transmit-receive RF surface coil, optimized to encompass the subjects’ head, was used for signal reception transmit and receive. Image localization and automated shimming were performed prior to data acquisition to ensure field homogeneity. Whole-brain fMRI data were acquired using a gradient-echo EPI sequence with the following parameters: echo time (TE) = 8 ms, repetition time (TR) = 1,500 ms, flip angle = 90°, voxel resolution = 0.99 × 0.99 × 0.93 mm, scan matrix = 64 × 64, and field of view (FOV) = 64 mm. The protocol included 54 coronal slices with a thickness of 1 mm, and 1,200 volumes were collected over a 30-min acquisition period. For anatomical reference and spatial normalization, distortion-matched structural images were acquired using a spin-echo EPI sequence with the following parameters: TE = 17.5 ms, TR = 1,500 ms, flip angle = 90°, 8 signal averages, scan matrix = 64 × 64, and FOV = 64 mm. The anatomical dataset consisted of 54 coronal slices (1 mm thickness), precisely aligned to match the functional imaging slices. Throughout MRI scanning sessions, subjects were continuously monitored in real-time via a live video feed (12 M camera, MRC Systems GmbH, Heidelberg, Germany). Physiological monitoring of heart rate and blood oxygen saturation (SpO₂) was performed using the Nonin 7500 pulse oximeter (Nonin Medical Inc., Plymouth, MN, USA), with measurements recorded at 5-min intervals to ensure subjects’ safety.

#### 2.4.3. MRI Data Preprocessing

Data were visually inspected for artifacts, and quality control was performed using the MRI Quality Control (MRIQC) tool [27]. Intensity spikes were identified and corrected using a custom program (https://github.com/bbfrederick/spikefix), with a threshold of 1 mm framewise displacement. Preprocessing was carried out using FMRIB’s Software Library (FSL, Oxford University, UK), including the removal of the first 10 volumes from each scan to allow for data stabilization. Head motion correction was performed using the MCFLIRT tool in FSL [28], with the resulting 12 motion correction parameters used as nuisance regressors.

Spatial smoothing was applied using a Gaussian kernel with a 2.0 mm full-width at half-maximum (FWHM), and temporal filtering was performed with a high-pass filter at a 100 s cutoff (0.01 Hz). Session-averaged functional volumes were aligned to the VALiDATe T2-weighted template [29] using a 12 degrees of freedom (DOF) affine transformation, followed by adjustment for nonlinear distortion fields using the JIP analysis toolkit (www.nitrc.org/projects/jip). Skull-stripping was applied to isolate brain tissue from non-brain structures.

#### 2.4.4. Analysis of Neural Responses to CS

A whole-brain analysis was conducted using FSL’s FEAT (FMRI Expert Analysis Tool) to examine blood oxygen level-dependent (BOLD) signal changes in response to CS presentation across the three experimental phases: Pre-C, Post-C, and Post-C+SCH. At the first (subject-level) analysis, event-related general linear models (GLMs) were constructed for each session, incorporating separate regressors for CS^+^ and CS^−^ trials. Each stimulus condition initially consisted of 23 trials; however, to enhance the reliability of cue-reactivity analyses, only trials exhibiting a stimulus-evoked BOLD response in the primary visual cortex during CS presentation were retained. This criterion ensured the inclusion of trials reflecting adequate visual engagement with the stimuli. Following this selection procedure, each subject averaged 13.44 ± 1.303 (SEM) trials per condition included in the final analysis. A separate regressor was defined for each session within each subject. Explanatory variables (EVs) were convolved with a gamma function (phase = 0, standard deviation = 3, mean lag = 6) to model the hemodynamic response, ensuring alignment with the expected temporal profile of the BOLD signal. To mitigate motion-related artifacts, subject-wise motion parameters (12 DOF) were included as nuisance regressors to account for residual motion effects. Contrast images were computed for four conditions: CS^+^ activation, CS^+^ deactivation, CS^-^ activation, and CS^-^ deactivation. At the second (group-level) analysis, a mixed-effects model (FLAME 1+2) was implemented to account for both within-and between-subject variance. A GLM-based ANOVA was conducted to compare contrast images across Pre-C, Post-C, and Post-C+SCH phases, evaluating phase-specific differences in neural responses to CS. Statistical inference was conducted using a voxel-wise threshold of Z > 3.1, with cluster-level family-wise error correction (cluster significance threshold, p < 0.05) based on Gaussian random field theory to control for multiple comparisons [30]. Brain regions involved in the identified significant clusters were delineated using a standardized squirrel monkey brain atlas [31] and independently verified by at least two experienced neuroscientists..

Time-series BOLD data were extracted from regions of interest (ROIs; masked as 3D spherical volumes with a 0.5 search radius) that exhibited significant CS^+^-evoked activation differences in both the Post-C > Pre-C and Post-C > Post-C+SCH contrasts, allowing for an analysis of the temporal dynamics of neural responses to the CS^+^ across experimental phases. Signal intensities were standardized across sessions using mean-based scaling to mitigate inter-session variability. Each stimulus presentation trial was segmented into an 18-sec epoch, consisting of a 6-sec pre-CS baseline, a 6-sec CS presentation period, and a 6-sec post-CS interval. To account for inter-trial variability, BOLD responses within each trial were normalized to the subjects’ baseline, defined as the mean BOLD signal intensity over the 6-sec pre-stimulus period. Normalized BOLD response was then calculated using the formula: ((BOLD signal / mean pre-stimulus BOLD signal) × 100) – 100, yielding the percentage signal change relative to the baseline. These normalized BOLD responses for each ROI were plotted separately for the three experimental phases: Pre-C, Post-C, and Post-C+SCH. To quantify stimulus-evoked neural activation, statistical analyses were conducted on the area under the curve (AUC) of the BOLD response within the 6–18-sec trial window.

#### 2.4.5. Analysis of Functional Connectivity between ROIs

Resting-state functional connectivity (rsFC) analysis was performed to evaluate the effects of SCH administration on brain network dynamics, specifically focusing on alterations in rsFC prior to CS exposure (480-sec baseline period). This approach was adopted to isolate and assess the effects of SCH administration, excluding any confounding influence from CS exposure. An ROI-based approach was employed, targeting key prefrontal subregions and the amygdala, as previous studies have demonstrated that rsFC within prefrontal-amygdala circuitry is a key determinant of fear response [32, 33]. Based on the VALiDATe29 Squirrel Monkey Brain Atlas [29], 10 ROIs were anatomically delineated and defined as 3D spherical masks (0.5 mm radius) centered on atlas-based coordinates: dmPFC, ventromedial prefrontal cortex (vmPFC), orbitofrontal cortex (OFC), left and right dorsolateral prefrontal cortex (dlPFC_L, dlPFC_R), left and right ventrolateral prefrontal cortex (vlPFC_L, vlPFC_R), left and right lateral orbitofrontal cortex (LOFC_L, LOFC_R), and the amygdala (Amyg). For each ROI, the baseline BOLD time-series (the first 480 secs) were extracted for analysis. Pairwise Pearson correlation coefficients were calculated across all ROI pairs using custom python scripts, resulting in a 10×10 connectivity matrix for each subject. To ensure normality and facilitate group-level comparisons, individual correlation coefficients were Fisher z-transformed prior to statistical analysis [34].

### 2.5. Correlation between Neural and Behavioral Responses to CS^+^

To assess whether CS+-evoked neural activation in specific brain regions was associated with behavioral performance, brain–behavior correlation analyses were conducted. For each subject and session, mean β-values were extracted from regions showing significant activation differences in both the Post-C > Pre-C and Post-C > Post-C+SCH contrasts. Behavioral performance was quantified as the response rate during CS+ presentations across experimental phases. Pearson correlation analyses were then performed to examine the relationship between region-specific neural activation and CS+-elicited behavioral responses.

### 2.6. Statistical Analysis

Statistical analyses were performed using Prism 9.0 software (GraphPad Software, Inc., San Diego, CA). For pairwise comparisons between two conditions (e.g., Pre-C vs. Post-C in Figure 1 and Post-C vs. Post-C+SCH in Figure 5), one-tailed paired t-tests were conducted to test directional hypotheses. Given the small sample size and the reduced statistical power associated with it, corrections for multiple comparisons were not applied to minimize the risk of Type II errors. For comparisons involving all three conditions, repeated-measures one-way analysis of variance (ANOVA) was employed, followed by Dunnett’s post hoc test for Figure 1 or Tukey’s post hoc test for Figure 3 to correct for multiple comparisons. Statistical significance thresholds were set at *p < 0.05, **p < 0.01, and ***p < 0.001.

**Figure 1.**
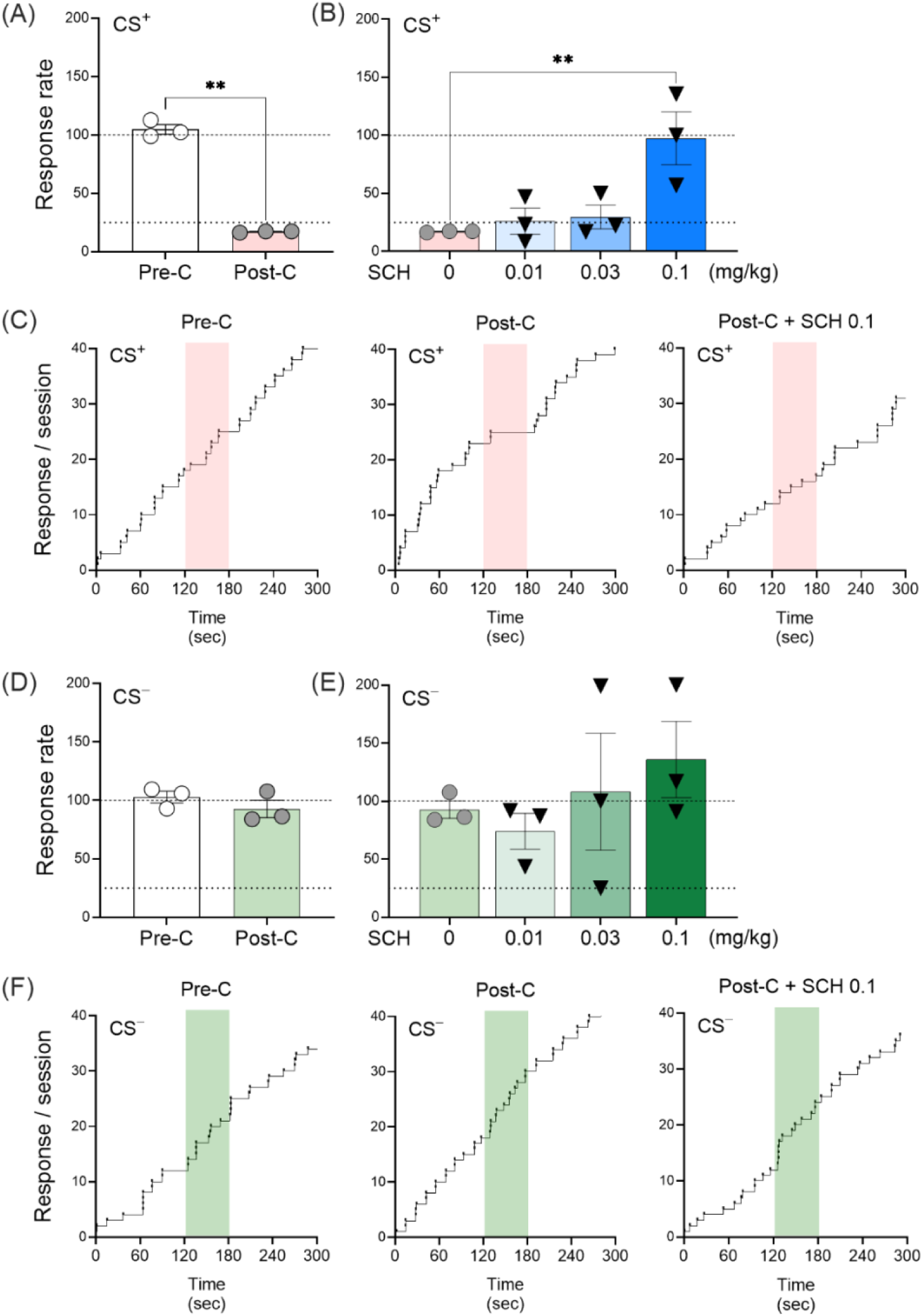
Changes in CS-associated behavioral responses across experimental phases. Subjects (n = 3) underwent a multi-phase conditioning paradigm consisting of operant training, in which the completion of an operant response was reinforced with milk delivery (0.3 mL/trial), followed by Pavlovian fear conditioning, where a CS^+^ was paired with an aversive US (0.5 mA electric stimulation), whereas the CS^−^ remained unpaired. During the test phase, behavioral responses to each CS were assessed by response rates, calculated as the ratio of the number of responses during the CS presentation period (120–180 sec; without US) to the number of responses during the baseline pre-CS interval (0–120 sec). (A) Response rate during CS^+^ presentation was significantly reduced in the Post-C phase compared to the Pre-C phase. (B) SCH administration (0.1 mg/kg) significantly attenuated CS^+^-induced suppression of operant responding. (C) Representative response patterns during CS^+^ sessions across experimental phases. (D) Response rate during CS^−^ presentation did not significantly differ between the Pre-C and Post-C phases. (E) No significant changes in CS^−^-associated behavioral responses were observed following SCH treatment at any dose. (F) Representative response patterns during CS^−^ sessions across experimental phases. Data are presented as mean ± SEM. Statistical significance shown as *p < 0.05, **p < 0.01.

**Figure 2.**
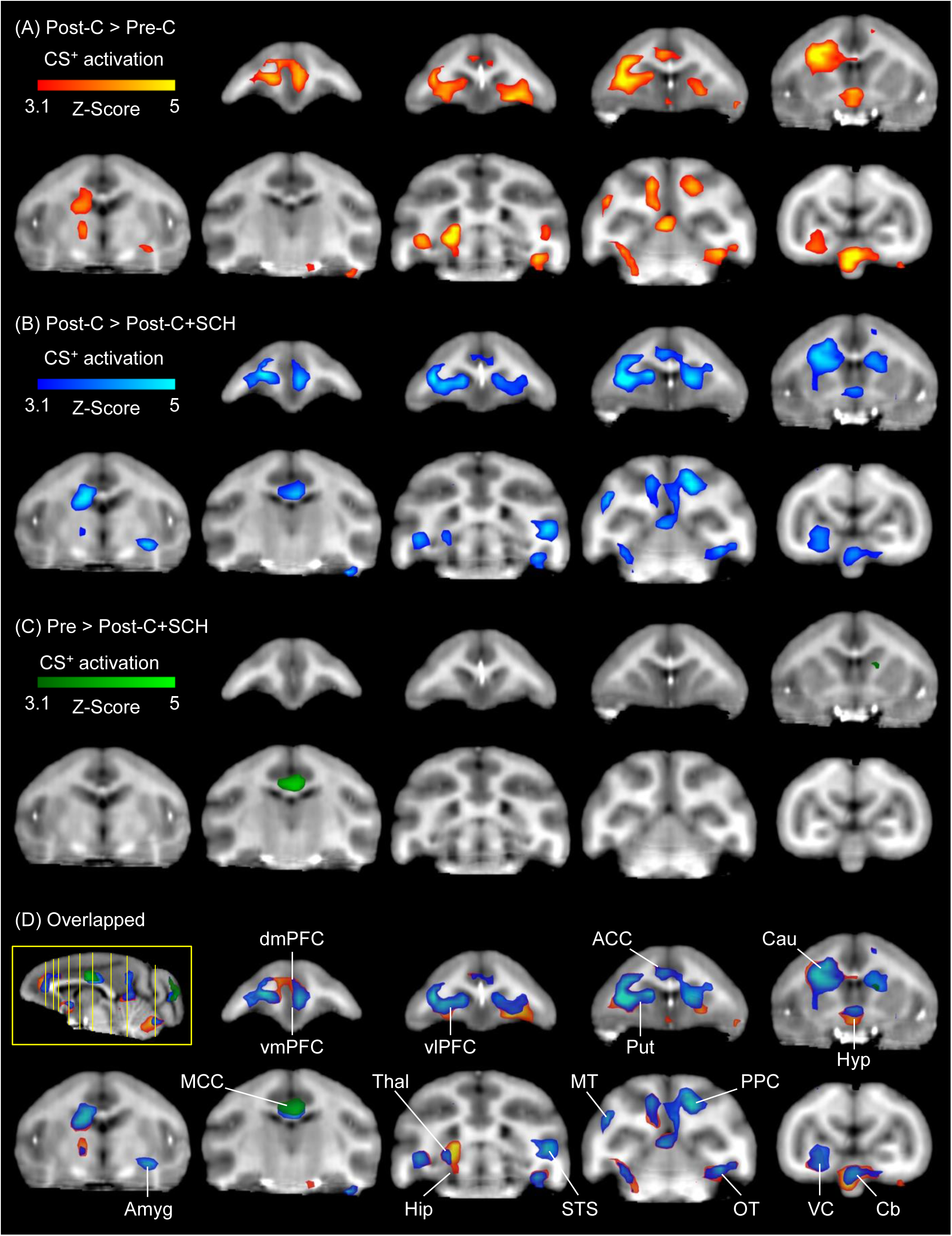
Whole-brain mapping of CS^+^-evoked BOLD response across experimental phases. Whole-brain voxel-wise analyses were conducted to assess differences in CS^+^-evoked BOLD responses across three experimental phases: Pre-C, Post-C, and Post-C+SCH. CS^+^-evoked activation is shown (A) during the Post-C phase compared to the Pre-C phase (Post-C > Pre-C). (B) following SCH administration in the Post-C+SCH phase relative to the Post-C phase (Post-C > Post-C+SCH) and, (C) between the Pre-C and Post-C+SCH phases (Pre-C > Post-C+SCH). (D) The sagittal image shows the locations of the displayed coronal slices along the anterior-posterior axis at the following coordinates: 18, 15.5, 14, 10.5, 7, 3, –3, –8, and –17. Overlapping spatial maps from contrasts (A)–(C) are visualized with semi-transparent overlays (opacity = 80%), revealing substantial spatial convergence between the Post-C > Pre-C and Post-C > Post-C+SCH clusters. Anatomical localizations of significant clusters were determined using a standardized squirrel monkey brain atlas [31]. Linear interpolation was applied using FSLeyes. Brain regions involved in the identified significant clusters included: dorsomedial prefrontal cortex (dmPFC), ventromedial prefrontal cortex (vmPFC), ventrolateral prefrontal cortex (vlPFC), anterior cingulate cortex (ACC), putamen (Put), caudate (Cau), hypothalamus (Hyp), amygdala (Amyg), middle cingulate cortex (MCC), thalamus (Thal), hippocampus (Hip), superior temporal association cortex (STS), middle temporal cortex (MT), posterior parietal cortex (PPC), occipito-temporal cortex (OT), visual cortex (VC), and cerebellum (Cb).

**Figure 3.**
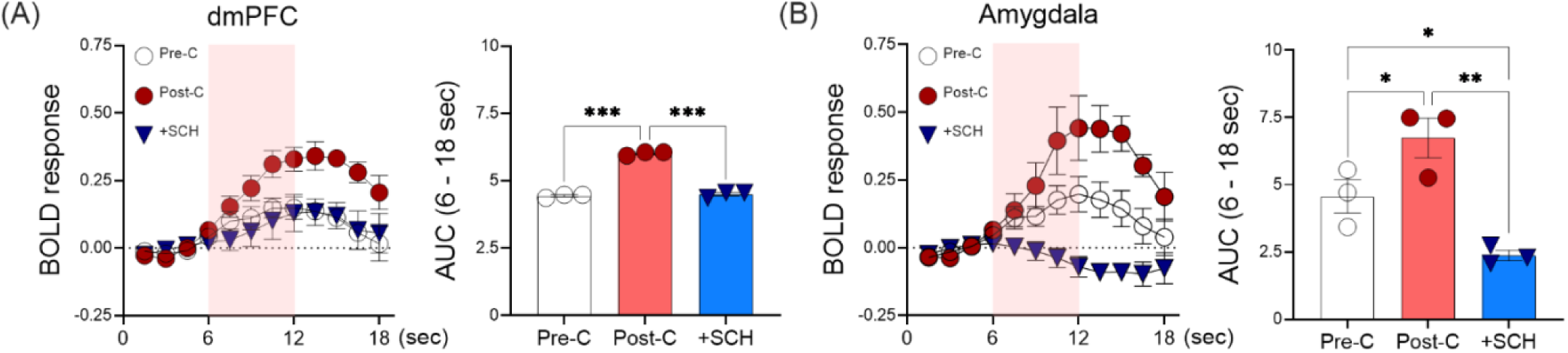
Changes in CS^+^-associated BOLD response within key ROIs across experimental phases. Time-series BOLD data were extracted from the dorsomedial prefrontal cortex (dmPFC) and amygdala (Amyg) to visualize CS^+^-evoked neural responses across experimental phases. Each CS trial was segmented into an 18-sec epoch comprising a pre-CS baseline period (0–6 sec), a CS presentation period (6–12 sec), and a post-CS interval (12–18 sec). Group-averaged BOLD response time courses (n = 3) were plotted for the three experimental phases: Pre-C, Post-C, and Post-C+SCH. To quantify CS^+^-evoked neural activation, the AUC of the BOLD response (%) from 6 to 18 secs was calculated for each phase. In the (A) dmPFC and (B) amygdala, CS^+^-evoked BOLD responses increased in the Post-C phase compared to the Pre-C phase. This conditioning-induced increase in BOLD responses was significantly attenuated following SCH administration (+SCH). AUC analyses revealed statistically significant differences across phases, consistent with these observations. Data are presented as mean ± SEM. Statistical significance was defined as *p < 0.05, **p < 0.01, ***p < 0.001.

**Figure 4.**
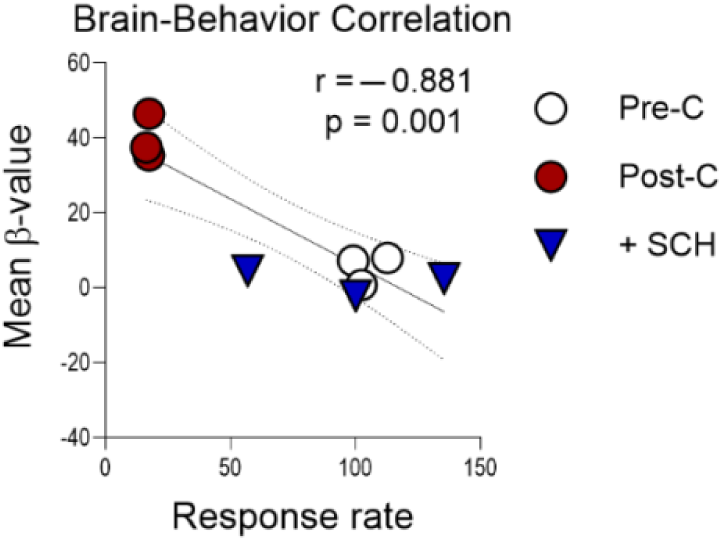
Negative correlation between CS^+^-evoked BOLD activation and behavioral response rates. For each subject (n = 3) and session, neural and behavioral responses to CS^+^ were quantified and included in a correlation analysis. Mean β-values were extracted from ROIs that showed significant activation differences in both the Post-C > Pre-C and Post-C > Post-C+SCH contrasts. Operant response rates during CS^+^ presentations were used as the behavioral measure. Pearson correlation analysis was conducted to assess the relationship between CS^+^-evoked neural activation and behavioral suppression, and the correlation coefficient (r) and statistical significance (p) were reported.

**Figure 5.**
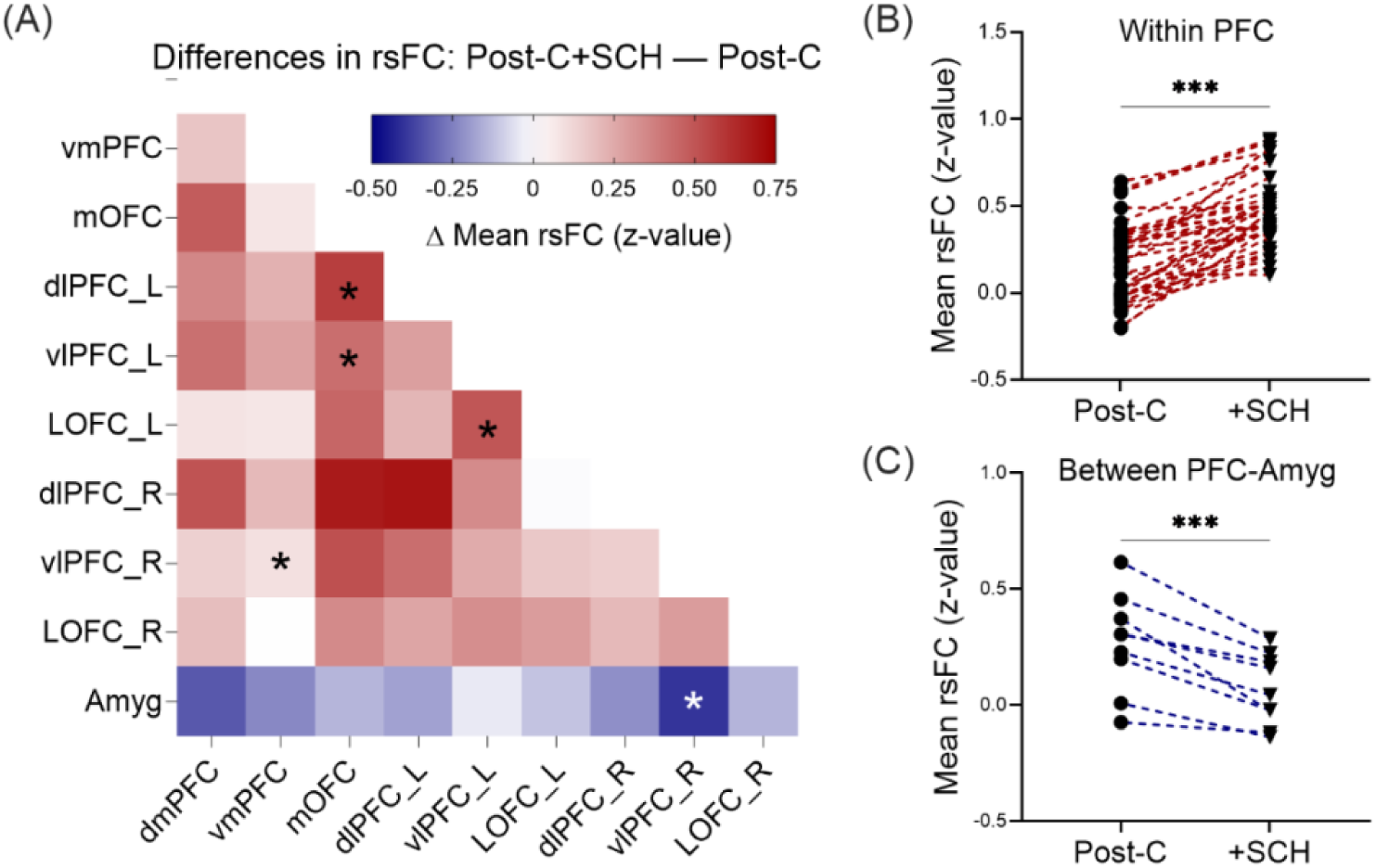
SCH-induced alterations in rsFC within prefrontal-amygdala circuitry. ROI-to-ROI connectivity analyses were conducted to assess changes in rsFC within the prefrontal-amygdala circuitry following SCH administration. Resting-state BOLD signals acquired during a 480-sec baseline period were used to estimate rsFC. Pearson’s correlation coefficients between ROI pairs were Fisher z-transformed, and group-level (n = 3) differences in mean rsFC between the Post-C and Post-C+SCH phases were evaluated. (A) The rsFC matrix across all ROIs shows that SCH administration generally increased rsFC within PFC subregions, while reducing rsFC between the amygdala and PFC subregions. (B) Mean rsFC within PFC subregions was significantly enhanced in the Post-C+SCH phase compared to the Post-C phase. (C) Mean rsFC between PFC subregions and the amygdala was significantly decreased in the Post-C+SCH phase relative to the Post-C phase. ROIs included: dorsomedial prefrontal cortex (dmPFC), ventromedial prefrontal cortex (vmPFC), orbitofrontal cortex (OFC), left and right dorsolateral prefrontal cortex (dlPFC_L, dlPFC_R), left and right ventrolateral prefrontal cortex (vlPFC_L, vlPFC_R), left and right lateral orbitofrontal cortex (OLFC_L, OLFC_R), and the amygdala (Amyg). Statistical significance was defined as *p < 0.05, ***p < 0.001.

## 3. Results

### 3.1. Effects of NOP Activation on Conditioned Behavioral Responses to CS

During the Pre-C phase, operant responding was stably maintained across the session regardless of CS^+^ or CS^−^ presentation (Figure 1C and 1F, left panels). However, in the Post-C phase, the response rate during CS^+^ presentations (Mean ± SEM, 17.03 ± 0.365) was significantly reduced compared to the Pre-C phase (104.7 ± 4.129), indicating CS^+^-induced suppression of operant responding (Figure 1A; t = 19.55, df = 2, p = 0.001). In contrast, responses during the Pre-CS and Post-CS periods did not significantly differ across experimental phases (Figure S1B), suggesting that the suppression of operant responding was both stimulus-specific and temporally constrained to the CS^+^ presentation period. Regarding the suppression of operant responding during CS^+^ presentation, administration of SCH produced a significant main effect of Dose (Figure 1B; F (3,6) = 8.853, p = 0.013), with the 0.1 mg/kg dose significantly attenuating conditioned suppression (97.43 ± 22.75; p = 0.009). In contrast, CS^−^ presentation did not induce significant changes in operant responding, regardless of the experimental phase, with no significant difference observed in the response rate between the Pre-C (102.8 ± 4.988) and Post-C (92.64 ± 7.556) phases during CS^−^ presentation (Figure 1D; t = 1.409, df = 2, p = 0.147). Furthermore, SCH treatment did not significantly alter the behavioral responses associated with CS^−^ exposure (Figure 1E; F = 0.579, p = 0.65). Figure 1C and 1F illustrate the response patterns during CS^+^ and CS^−^ sessions, respectively, across the three experimental phases (Pre-C, Post-C, and Post-C+SCH 0.1) in a representative subject, with the response patterns for all subjects provided in Figure S2.

### 3.2. Effects of NOP Activation on Neural Responses to CS After Conditioning

Whole-brain voxel-wise analyses revealed significant differences in CS-evoked BOLD responses across three experimental phases: Pre-C, Post-C, and Post-C+SCH (Table 1). Figure 2A shows that CS+-evoked activation significantly increased during the Post-C phase compared to the Pre-C phase (Post-C > Pre-C), particularly in regions such as the dmPFC, vmPFC, bilateral vlPFC, and amygdala, (see complete list of regions in Table 1). Following SCH administration, a significant decrease in CS^+^-evoked activation was observed during the Post-C+SCH phase relative to the Post-C phase (Post-C > Post-C+SCH; Figure 2B), primarily in regions overlapping with the Post-C > Pre-C contrast, as shown in Figure 2D, along with additional reduction in the midcingulate cortex (MCC). Figure 2C demonstrates that SCH administration attenuated CS^+^-evoked activation in the MCC, as evidenced by significantly greater activation during the Pre-C phase compared to the Post-C+SCH phase (Pre-C > Post-C+SCH).

**Table 1.**
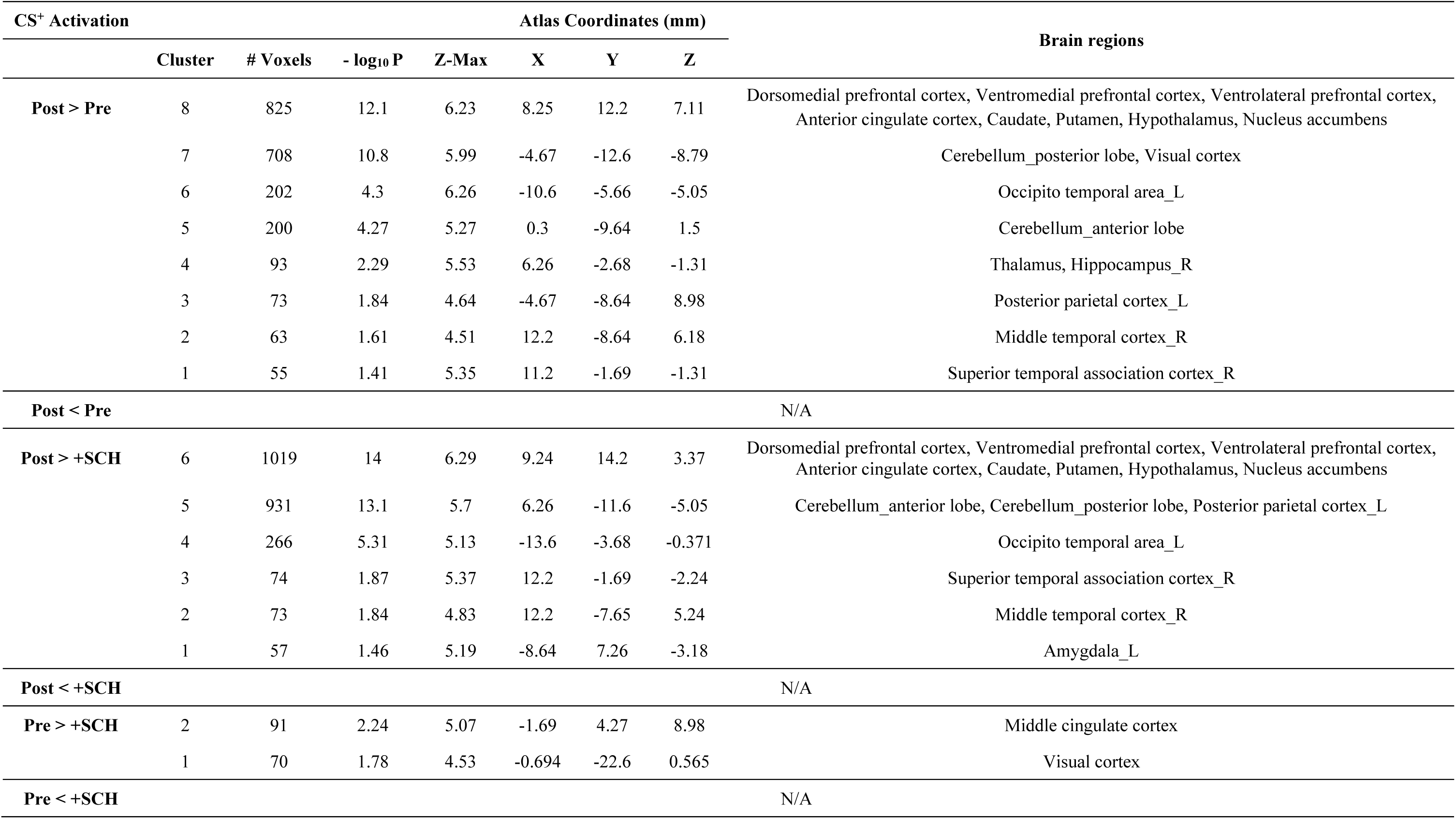
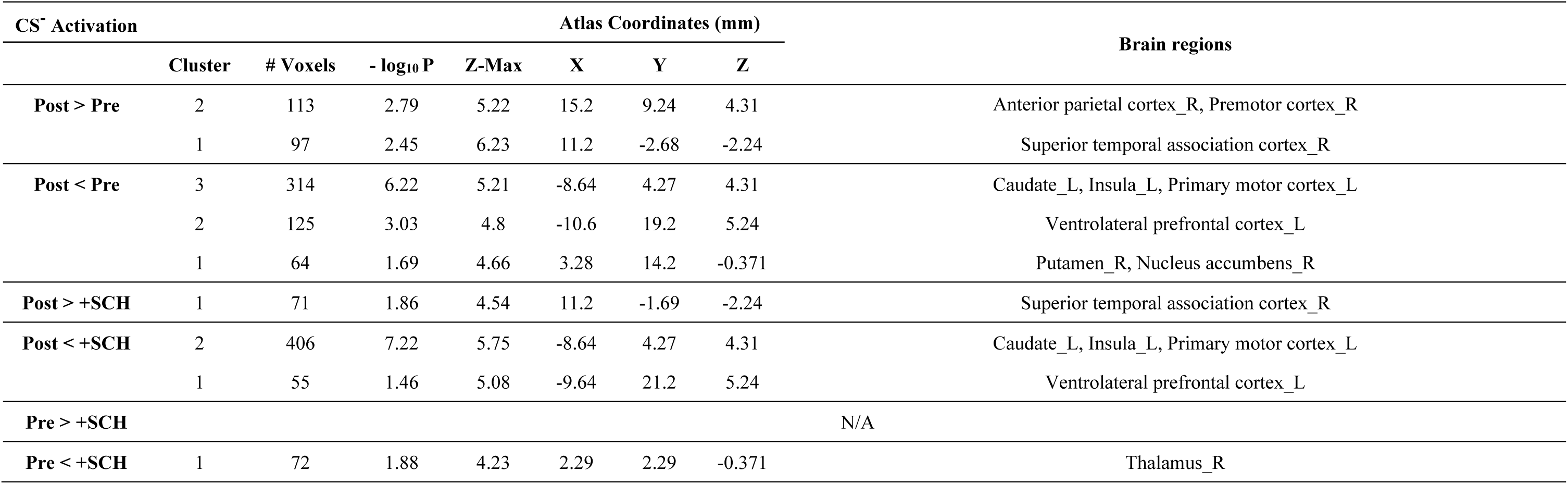
Brain regions exhibiting significant differences in CS-evoked BOLD responses across the three experimental phases (Pre-C, Post-C, and Post-C+SCH). Whole-brain voxel-wise analyses were performed using a GLM-based repeated-measures ANOVA to identify phase-dependent alterations in neural responses to each CS (CS^+^ or CS^-^). Statistical significance was determined using a voxel-level Z-threshold of 3.1 and cluster-level correction for multiple comparisons based on FWE correction (p < 0.05). The brain regions encompassed within significant clusters were anatomically delineated using a standardized squirrel monkey brain atlas. R = right; L = left.

| CS<sup>+</sup> Activation |  | Atlas Coordinates (mm) |  |  |  |  |  | Brain regions |
| --- | --- | --- | --- | --- | --- | --- | --- | --- |
|  | Cluster | # Voxels | - log<sub>10</sub> P | Z-Max | X | Y | Z |  |
| Post > Pre | 8 | 825 | 12.1 | 6.23 | 8.25 | 12.2 | 7.11 | Dorsomedial prefrontal cortex, Ventromedial prefrontal cortex, Ventrolateral prefrontal cortex, Anterior cingulate cortex, Caudate, Putamen, Hypothalamus, Nucleus accumbens |
|  | 7 | 708 | 10.8 | 5.99 | -4.67 | -12.6 | -8.79 | Cerebellum_posterior lobe, Visual cortex |
|  | 6 | 202 | 4.3 | 6.26 | -10.6 | -5.66 | -5.05 | Occipito temporal area_L |
|  | 5 | 200 | 4.27 | 5.27 | 0.3 | -9.64 | 1.5 | Cerebellum_anterior lobe |
|  | 4 | 93 | 2.29 | 5.53 | 6.26 | -2.68 | -1.31 | Thalamus, Hippocampus_R |
|  | 3 | 73 | 1.84 | 4.64 | -4.67 | -8.64 | 8.98 | Posterior parietal cortex_L |
|  | 2 | 63 | 1.61 | 4.51 | 12.2 | -8.64 | 6.18 | Middle temporal cortex_R |
|  | 1 | 55 | 1.41 | 5.35 | 11.2 | -1.69 | -1.31 | Superior temporal association cortex_R |
| Post < Pre |  |  |  |  |  |  |  | N/A |
| Post > +SCH | 6 | 1019 | 14 | 6.29 | 9.24 | 14.2 | 3.37 | Dorsomedial prefrontal cortex, Ventromedial prefrontal cortex, Ventrolateral prefrontal cortex, Anterior cingulate cortex, Caudate, Putamen, Hypothalamus, Nucleus accumbens |
|  | 5 | 931 | 13.1 | 5.7 | 6.26 | -11.6 | -5.05 | Cerebellum_anterior lobe, Cerebellum_posterior lobe, Posterior parietal cortex_L |
|  | 4 | 266 | 5.31 | 5.13 | -13.6 | -3.68 | -0.371 | Occipito temporal area_L |
|  | 3 | 74 | 1.87 | 5.37 | 12.2 | -1.69 | -2.24 | Superior temporal association cortex_R |
|  | 2 | 73 | 1.84 | 4.83 | 12.2 | -7.65 | 5.24 | Middle temporal cortex_R |
|  | 1 | 57 | 1.46 | 5.19 | -8.64 | 7.26 | -3.18 | Amygdala_L |
| Post < +SCH |  |  |  |  |  |  |  | N/A |
| Pre > +SCH | 2 | 91 | 2.24 | 5.07 | -1.69 | 4.27 | 8.98 | Middle cingulate cortex |
|  | 1 | 70 | 1.78 | 4.53 | -0.694 | -22.6 | 0.565 | Visual cortex |
| Pre < +SCH |  |  |  |  |  |  |  | N/A |

| CS<sup>-</sup> Activation |  | Atlas Coordinates (mm) |  |  |  |  |  | Brain regions |
| --- | --- | --- | --- | --- | --- | --- | --- | --- |
|  | Cluster | # Voxels | - log<sub>10</sub> P | Z-Max | X | Y | Z |  |
| Post > Pre | 2 | 113 | 2.79 | 5.22 | 15.2 | 9.24 | 4.31 | Anterior parietal cortex_R, Premotor cortex_R |
|  | 1 | 97 | 2.45 | 6.23 | 11.2 | -2.68 | -2.24 | Superior temporal association cortex_R |
| Post < Pre | 3 | 314 | 6.22 | 5.21 | -8.64 | 4.27 | 4.31 | Caudate_L, Insula_L, Primary motor cortex_L |
|  | 2 | 125 | 3.03 | 4.8 | -10.6 | 19.2 | 5.24 | Ventrolateral prefrontal cortex_L |
|  | 1 | 64 | 1.69 | 4.66 | 3.28 | 14.2 | -0.371 | Putamen_R, Nucleus accumbens_R |
| Post > +SCH | 1 | 71 | 1.86 | 4.54 | 11.2 | -1.69 | -2.24 | Superior temporal association cortex_R |
| Post < +SCH | 2 | 406 | 7.22 | 5.75 | -8.64 | 4.27 | 4.31 | Caudate_L, Insula_L, Primary motor cortex_L |
|  | 1 | 55 | 1.46 | 5.08 | -9.64 | 21.2 | 5.24 | Ventrolateral prefrontal cortex_L |
| Pre > +SCH |  | N/A |  |  |  |  |  |  |
| Pre < +SCH | 1 | 72 | 1.88 | 4.23 | 2.29 | 2.29 | -0.371 | Thalamus_R |

Time-series analyses were performed on ROIs identified in Table 1, which exhibited significant increase in CS^+^-evoked activation following fear conditioning. Across all ROIs, BOLD response to CS^+^ increased during the Post-C phase relative to Pre-C and was subsequently attenuated following SCH administration. Representative time-series data from two ROIs, including the dmPFC and the amygdala, are shown in Figure 3. In the dmPFC, a significant main effect of experimental phase was observed (Figure 3A; F = 37.1, p < 0.001), characterized by a robust increase in the area under the curve (AUC) from Pre-C to Post-C (p < 0.001), followed by a significant reduction from Post-C to Post-C+SCH (p < 0.001). Likewise, the amygdala exhibited a significant main effect of phase (Figure 3B; F = 31.75, p = 0.004), with post hoc comparisons indicating significant differences across all phase contrasts: increased AUC from Pre-C to Post-C (p = 0.036), and decreased AUC from Post-C to Post-C+SCH (p = 0.003) as well as from Pre-C to Post-C+SCH (p = 0.034).

### 3.3. Relation Between CS^+^-Evoked BOLD Activation and Conditioned Behavioral Suppression

A significant negative correlation was observed between mean β-values from the defined ROIs and response rates during CS^+^ presentation (Figure 4; r =-0.881, p = 0.001). Specifically, the Post-C phase exhibited significantly elevated β-values concomitant with reduced response rates relative to the Pre-C and Post-C+SCH phases, indicating that increased neural activation within these ROIs was associated with conditioned behavioral suppression.

### 3.4. Effects of NOP Agonism on rsFC of the Prefrontal-Amygdala Circuit

Administration of SCH induced significant alterations in prefrontal–amygdala network dynamics (Figure 5A). Notably, increased rsFC was observed among several PFC subregions, including vmPFC–vlPFC_R (p = 0.014, t = 5.886), mOFC–dlPFC_L (p = 0.013, t = 6.090), mOFC–vlPFC_L (p = 0.041, t = 3.251), and LOFC_L–vlPFC_L (p = 0.027, t = 4.095). In contrast, a significant reduction in rsFC was detected between vlPFC_R and the amygdala (p = 0.027, t = 4.120). Group-level analyses further corroborated these findings, demonstrating a robust enhancement of intra-PFC connectivity (Figure 5B; p < 0.001, t = 9.930, df = 35), along with a significant decrease in connectivity between PFC and amygdala (Figure 5C; p < 0.001, t = 5.613, df = 8).

## 4. Discussion

The present study demonstrates that activation of the NOP system via systemic administration of SCH attenuates behavioral responses to aversive stimuli and modulates neural activity and functional connectivity within fear-associated circuits in NHPs. From a methodological perspective, this study builds upon existing translational approaches by employing a Pavlovian conditioning paradigm in combination with event-related fMRI [35], a technique that remains relatively underutilized in preclinical research. This methodological convergence enhances cross-species translational validity and enables a more comprehensive characterization of the neural substrates underlying maladaptive fear processing, thereby contributing to the development of mechanism-based therapeutic strategies for anxiety disorders.

In behavioral assessments, subjects exhibited robust suppression of operant responding during CS^+^ presentation following fear conditioning, indicative of conditioned behavioral inhibition elicited by an aversive stimulus. Notably, activation of the NOP receptor significantly attenuated this behavioral suppression, leading to a recovery of operant responding during CS^+^ presentation, consistent with prior preclinical findings that the NOP receptor agonist exerts anxiolytic-like effects in rodents [8, 9]. Although one potential interpretation of the restoration is that NOP receptor activation enhances general operant behavior, this explanation seems unlikely, as response during the pre-CS period were not elevated and were instead modestly reduced following NOP agonist administration (Figure S1C). These findings suggest that the primary effect of NOP receptor activation is not the enhancement of general motivated behavior, but rather the selective attenuation of conditioned fear-related behavioral responses to aversive stimuli. The specificity of this anxiolytic effect was further supported by the absence of significant behavioral effects of NOP activation during CS^−^ sessions, indicating that NOP receptor modulation selectively targets fear-associated behavioral inhibition without producing nonspecific effects on motor function or motivation.

Whole-brain voxel-wise analyses revealed robust BOLD activation in response to CS^+^ following fear conditioning across multiple neural networks, including regions implicated in fear processing [36–38] (amygdala, hypothalamus, caudate, putamen), fear expression and regulation [39–41] (dmPFC, vmPFC, vlPFC, ACC), and sensory integration [42–44] (visual cortex, parietal cortex, temporal cortex, thalamus, cerebellum). These findings align with previous studies demonstrating the engagement of fronto-limbic and sensorimotor networks in visually mediated fear signaling pathways [45]. Notably, SCH administration significantly attenuated CS^+^-evoked activation across most of these regions, indicating that NOP receptor activation suppresses conditioned neural responses to aversive stimuli. In addition, response to the CS^+^ in the MCC, a region implicated in threat detection and appraisal [46], was significantly reduced by SCH treatment relative to both the Pre-C and Post-C conditions (Figure 2B and C). A similar dampening effects on response to CS^−^ presentation were also observed. Specifically, during the Post-C phase, CS^−^ presentation evoked reduced neural activation in several fear-related regions (caudate, putamen, vlPFC), while increased activity was observed in sensory processing areas (temporal cortex) (Table 1). Taken together, NOP agonism blunted responses to both CS^+^ and CS^−^, suggesting that NOP signaling may diminish neural discrimination between aversive and non-aversive cues by modulating circuits involved in fear processing and sensory salience. This widespread, yet regionally selective modulatory effect likely reflects the neuroanatomical distribution of NOP receptors, which are densely expressed across key regions of the fronto-limbic circuitry, including the PFC, cingulate cortex, amygdala, hypothalamus, and dorsal striatum [7]. Further, considering that visual responses to fearful stimuli are modulated by the limbic regions including amygdala [47], the observed reduction in activity across sensory integration areas may suggest that NOP receptor agonism attenuates sensory processing of aversive stimuli, potentially as a downstream consequence of reduced limbic activation.

Time-series analysis was performed to examine how neural responses evolved over the course of CS presentations and to assess whether NOP activation modulated the transient or sustained effects of CS. This analysis provided superior temporal resolution compared to conventional block-averaged methods, enabling the detection of dynamic fluctuations in BOLD signals in response to CS presentations. Furthermore, this approach enabled the precise characterization of phasic changes in BOLD responses to CS^+^ across experimental phases, including the conditioning-induced increase and its subsequent attenuation following SCH administration, across all ROIs identified in the whole-brain analysis. Notably, representative regions such as the dmPFC and amygdala exhibited robust increases in CS^+^-evoked activation following conditioning, consistent with previous studies showing the activation in these regions during fear processing [48, 49]. The attenuation of this activation through NOP receptor agonism indicates that the NOP system serves as a key neuromodulatory pathway in regulating activity within fear-associated neural circuits.

Correlation analyses further supported the finding that increased activation within fear-associated ROIs was associated with decreased operant responding during CS^+^ presentations, particularly in the Post-C phase. Activation of the NOP receptor attenuated this heightened neural reactivity within fronto-limbic circuits in response to conditioned aversive stimuli, which was associated with a corresponding reduction in behavioral suppression, thereby providing mechanistic insight into the anxiolytic-like effects of NOP receptor agonism.

While NOP activation attenuated the response to the CS^+^, we also examined whether it influenced brain activity independently of stimulus presentation, specifically by modulating baseline functional connectivity prior to CS onset. rsFC analyses within the PFC-amygdala circuitry revealed bidirectional modulatory effects of NOP receptor activation. Contrary to the initial expectation that NOP receptor activation, similar to classical anxiolytics, would enhance top-down regulatory control by increasing connectivity between the PFC and amygdala [50], a significant decrease in functional connectivity between these regions was observed. Given that the PFC-amygdala pathway plays a central role in transmitting aversive stimulus information necessary for the initiation and expression of conditioned fear responses [51], this reduction may suggest that NOP receptor activation disrupts the transmission of fear-related signals, thereby attenuating bottom-up signaling. This finding aligns with prior studies demonstrating that reduced PFC-amygdala coupling is associated with attenuated fear expression, as evidenced by reduced behavioral reactivity to aversive stimuli [32]. Conversely, NOP receptor activation enhanced intra-PFC connectivity, indicating increased functional integration among PFC subregions involved in the regulation of affective responses [52]. This finding is in line with existing literature reporting that decreased intra-PFC connectivity is often observed in heightened fear-related anxiety conditions [53]. Taken together, these findings suggest that NOP receptor activation suppresses the bottom-up propagation of fear-related signals originating from the amygdala while concurrently strengthening intra-PFC coordination that facilitates regulatory control. This bidirectional mechanism may underlie the observed attenuation of both behavioral and neural responses to conditioned aversive stimuli.

Several considerations, particularly the relatively small sample size, which is a common constraint in nonhuman primate research, especially when using complex behavioral tasks, warrant careful consideration in the interpretation of these findings. In the rsFC analyses, although the importance of correcting for multiple comparisons to control for Type I error is well-established, the limited sample size substantially reduced statistical power, thereby increasing the likelihood of Type II error following correction [54]. Consequently, uncorrected results are presented with appropriate caution and transparency. Despite this limitation, the convergence of results across multiple methodological approaches, including behavioral paradigms, event-related BOLD responses, and rsFC analyses, supports the internal consistency and construct validity of the observed effects. Future studies that include larger sample sizes will be essential for enhancing statistical sensitivity and confirming the replicability of these findings.

In conclusion, the present study provides compelling evidence that pharmacological activation of the NOP receptor attenuates behavioral and neural responses to conditioned aversive stimuli in nonhuman primates. Specifically, NOP receptor agonism appeared to modulate functional interactions within the PFC-amygdala circuitry and suppresses aversive stimulus-evoked neural activation across the fronto-limbic network, thereby resulting in a selective attenuation of fear-related behavioral responses. Taken together, these findings suggest that NOP receptor agonism may be a promising therapeutic strategy for fear-related anxiety disorders.

## Acknowledgments

This research was supported by P50MH119467-04 grant from NIMH (KHH and DAP). The content is solely the responsibility of the authors and does not necessarily represent the official views of the National Institutes of Health. The authors thank Julia Cunningham for expert technical assistance in conducting experiments.

## Authorship

Kwang-Hyun Hur: Conceptualization, Investigation, Methodology, Formal analysis, Writing - Original Draft, Visualization, Funding acquisition

Diego A. Pizzagalli: Writing - Review & Editing, Supervision, Funding acquisition Jessi Stover: Investigation

Kenroy Cayetano: Software, Resources

Stephen J. Kohut: Conceptualization, Methodology, Resources, Writing - Review & Editing, Supervision

## Data Availability Statement

All data needed to evaluate the conclusions in the paper are present in the paper. Raw data from this study are available from the corresponding author upon reasonable request.

---

**Figure S1.**
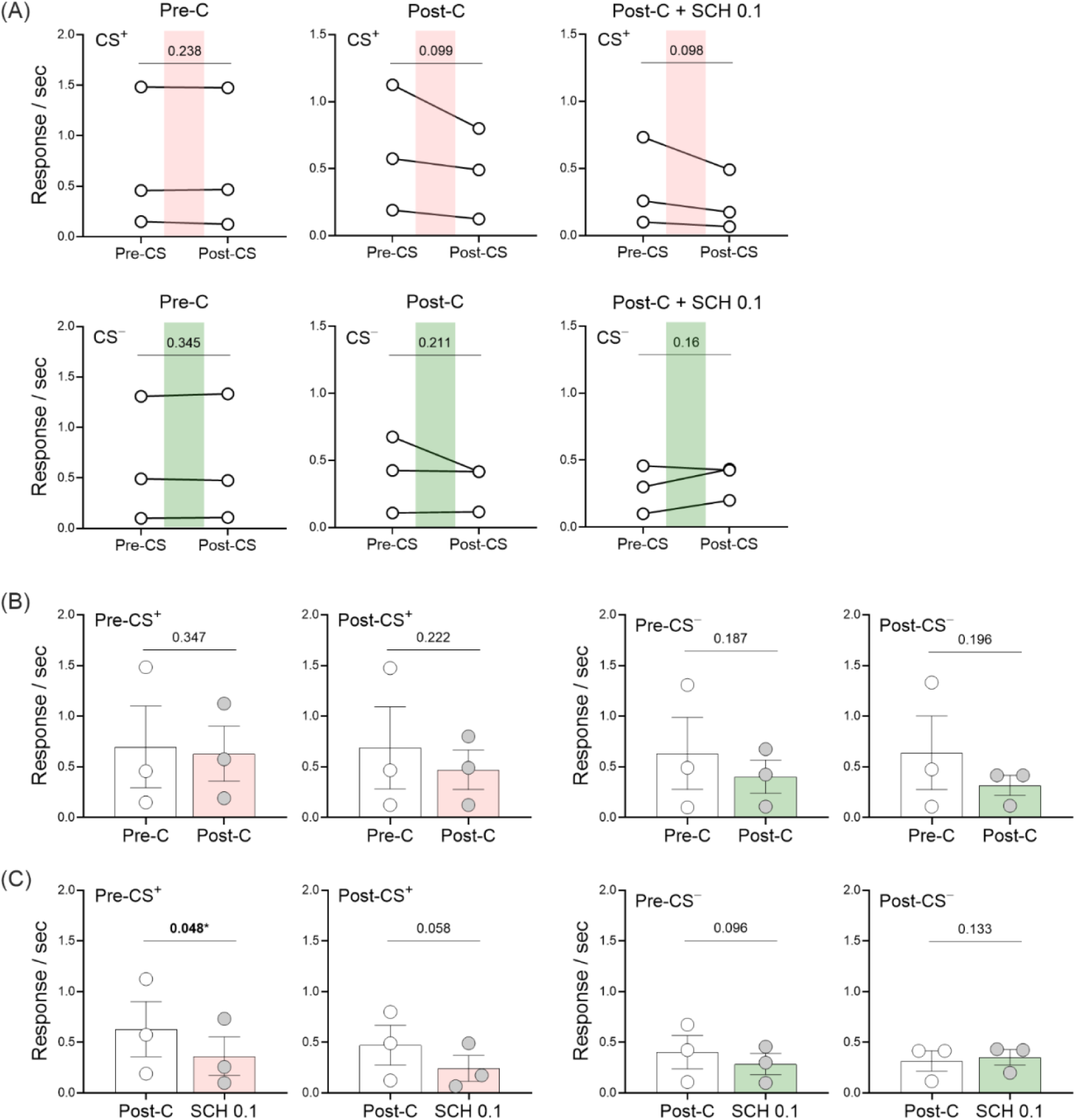
Comparison of responses during Pre-CS and Post-CS periods within and across experimental phases. (A) Responses during the Pre-CS and Post-CS periods were compared within each experimental phase (Pre-C, Post-C, and Post-C+SCH 0.1) for both CS^+^ and CS^−^ sessions. (B–C) Responses during each time window (Pre-CS and Post-CS) were compared across experimental phases: (B) between the Pre-C and Post-C phases, and (C) between the Post-C and Post-C+SCH 0.1 phases, in both CS+ and CS− sessions. Statistical significance was assessed using p-values and is indicated in the corresponding panels.

**Figure S2.**
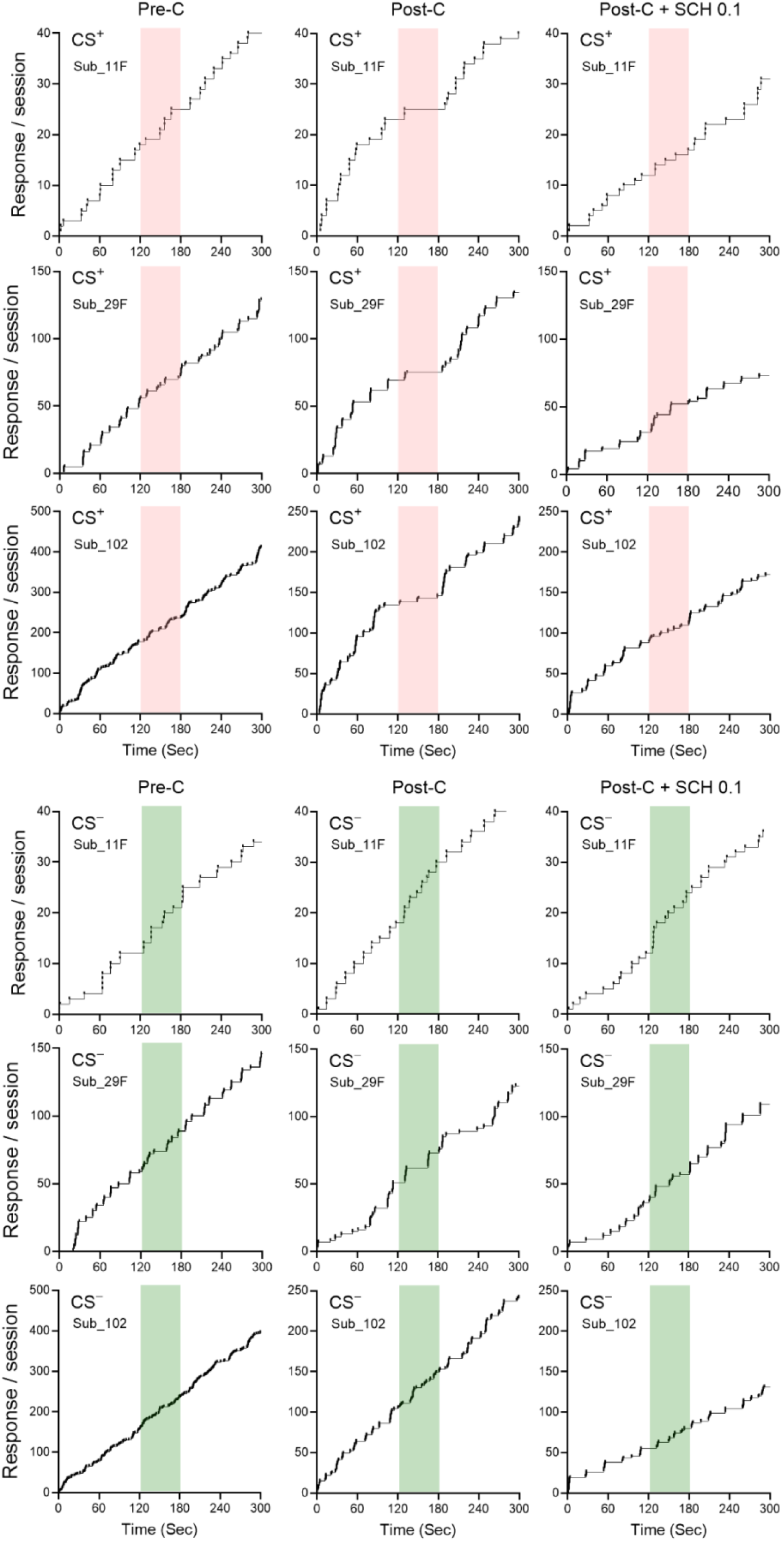
Response patterns during CS^+^ and CS^−^ sessions across the three experimental phases in all subjects.

## References

1. World Health Organization, Anxiety disorders. https://www.who.int/news-room/fact-sheets/detail/anxiety-disorders, 2023.

2. Hartley, C.A. and B. Casey, Risk for anxiety and implications for treatment: developmental, environmental, and genetic factors governing fear regulation. Annals of the New York Academy of Sciences, 2013. 1304(1): p. 1–13.

3. Shin, L.M. and I. Liberzon, The neurocircuitry of fear, stress, and anxiety disorders. Neuropsychopharmacology, 2010. 35(1): p. 169–191.

4. Duval, E.R., A. Javanbakht, and I. Liberzon, Neural circuits in anxiety and stress disorders: a focused review. Therapeutics and clinical risk management, 2015: p. 115–126.

5. Bystritsky, A., Treatment-resistant anxiety disorders. Molecular psychiatry, 2006. 11(9): p. 805–814.

6. Mallimo, E.M. and A.W. Kusnecov, The role of orphanin FQ/nociceptin in neuroplasticity: relationship to stress, anxiety and neuroinflammation. Frontiers in cellular neuroscience, 2013. 7: p. 173.

7. Witta, J., et al., Distribution of nociceptin/orphanin FQ in adult human brain. Brain research, 2004. 997(1): p. 24–29.

8. Ross, T.M., et al., A selective small molecule NOP (ORL-1 receptor) partial agonist for the treatment of anxiety. Bioorganic & Medicinal Chemistry Letters, 2015. 25(3): p. 602–606.

9. Jenck, F., et al., A synthetic agonist at the orphanin FQ/nociceptin receptor ORL1: anxiolytic profile in the rat. Proceedings of the National Academy of Sciences, 2000. 97(9): p. 4938–4943.

10. Gold, A.L., et al., Age differences in the neural correlates of anxiety disorders: An fMRI study of response to learned threat. American Journal of Psychiatry, 2020. 177(5): p. 454–463.

11. Britton, J.C., et al., Response to learned threat: An fMRI study in adolescent and adult anxiety. American Journal of Psychiatry, 2013. 170(10): p. 1195–1204.

12. McCabe, C., et al., Diminished neural processing of aversive and rewarding stimuli during selective serotonin reuptake inhibitor treatment. Biological psychiatry, 2010. 67(5): p. 439–445.

13. Qiao, N., et al., Update on Nonhuman Primate Models of Brain Disease and Related Research Tools. Biomedicines, 2023. 11(9): p. 2516.

14. Bridge, K., et al., Autoradiographic localization of 125I [Tyr14] nociceptin/orphanin FQ binding sites in macaque primate CNS. Neuroscience, 2003. 118(2): p. 513–523.

15. Estes, W.K. and B.F. Skinner, Some quantitative properties of anxiety. Journal of Experimental Psychology, 1941. 29(5): p. 390.

16. Park, J. and B. Moghaddam, Impact of anxiety on prefrontal cortex encoding of cognitive flexibility. Neuroscience, 2017. 345: p. 193–202.

17. Adewale, A.S., D.M. Platt, and R.D. Spealman, Pharmacological stimulation of group ii metabotropic glutamate receptors reduces cocaine self-administration and cocaine-induced reinstatement of drug seeking in squirrel monkeys. Journal of Pharmacology and Experimental Therapeutics, 2006. 318(2): p. 922–931.

18. Withey, S.L., et al., Fentanyl-induced changes in brain activity in awake nonhuman primates at 9.4 Tesla. Brain Imaging and Behavior, 2022. 16(4): p. 1684–1694.

19. Lamb, G.D., Understanding“within” versus“between” ANOVA Designs: Benefits and Requirements of Repeated Measures. 2003.

20. Kangas, B.D. and J. Bergman, Operant nociception in nonhuman primates. PAIN®, 2014. 155(9): p. 1821–1828.

21. Flynn, S.M., et al., Effects of stimulation of mu opioid and nociceptin/orphanin FQ peptide (NOP) receptors on alcohol drinking in rhesus monkeys. Neuropsychopharmacology, 2019. 44(8): p. 1476–1484.

22. Atack, J.R., et al., TPA023 [7-(1, 1-dimethylethyl)-6-(2-ethyl-2 H-1, 2, 4-triazol-3-ylmethoxy)-3-(2-fluorophenyl)-1, 2, 4-triazolo [4, 3-b] pyridazine], an agonist selective for α2-and α3-containing GABAA receptors, is a nonsedating anxiolytic in rodents and primates. The Journal of pharmacology and experimental therapeutics, 2006. 316(1): p. 410–422.

23. Yassin, W., et al., Resting-state networks of awake adolescent and adult squirrel monkeys using ultra-high field (9.4 T) functional magnetic resonance imaging. Eneuro, 2024. 11(5).

24. Schlauch, R.C., et al., Psychometric evaluation of the Substance Use Risk Profile Scale (SURPS) in an inpatient sample of substance users using cue-reactivity methodology. Journal of psychopathology and behavioral assessment, 2015. 37: p. 231–246.

25. Katwal, S.B., et al., Measuring relative timings of brain activities using fMRI. NeuroImage, 2013. 66: p. 436–448.

26. Kim, J.H., et al., Dynamics of the cerebral blood flow response to brief neural activity in human visual cortex. Journal of Cerebral Blood Flow & Metabolism, 2020. 40(9): p. 1823–1837.

27. Esteban, O., et al., MRIQC: Advancing the automatic prediction of image quality in MRI from unseen sites. PloS one, 2017. 12(9): p. e0184661.

28. Jenkinson, M., et al., Improved optimization for the robust and accurate linear registration and motion correction of brain images. Neuroimage, 2002. 17(2): p. 825–841.

29. Schilling, K.G., et al., The VALiDATe29 MRI based multi-channel atlas of the squirrel monkey brain. Neuroinformatics, 2017. 15: p. 321–331.

30. Worsley, K.J., Statistical analysis of activation images. Functional MRI: An introduction to methods, 2001. 14(1): p. 251–270.

31. Gergen, J.A. and P.D. MacLean, A stereotaxic atlas of the squirrel monkey’s brain (Saimiri sciureus). 1962: US Department of Health, Education, and Welfare, Public Health Service….

32. Ganella, D.E., et al., Prefrontal-amygdala connectivity and state anxiety during fear extinction recall in adolescents. Frontiers in human neuroscience, 2017. 11: p. 587.

33. Feng, P., et al., Memory consolidation of fear conditioning: bi-stable amygdala connectivity with dorsal anterior cingulate and medial prefrontal cortex. Social cognitive and affective neuroscience, 2014. 9(11): p. 1730–1737.

34. Van Dijk, K.R., et al., Intrinsic functional connectivity as a tool for human connectomics: theory, properties, and optimization. Journal of neurophysiology, 2010. 103(1): p. 297–321.

35. Reinhardt, I., et al., Neural correlates of aversive conditioning: development of a functional imaging paradigm for the investigation of anxiety disorders. European Archives of Psychiatry and Clinical Neuroscience, 2010. 260: p. 443–453.

36. Öhman, A., The role of the amygdala in human fear: automatic detection of threat. Psychoneuroendocrinology, 2005. 30(10): p. 953–958.

37. Gross, C.T. and N.S. Canteras, The many paths to fear. Nature Reviews Neuroscience, 2012. 13(9): p. 651–658.

38. Ferreira, T., et al., The indirect amygdala–dorsal striatum pathway mediates conditioned freezing: Insights on emotional memory networks. Neuroscience, 2008. 153(1): p. 84–94.

39. Milad, M.R., et al., A role for the human dorsal anterior cingulate cortex in fear expression. Biological psychiatry, 2007. 62(10): p. 1191–1194.

40. Vidal-Gonzalez, I., et al., Microstimulation reveals opposing influences of prelimbic and infralimbic cortex on the expression of conditioned fear. Learning & memory, 2006. 13(6): p. 728–733.

41. Sotres-Bayon, F. and G.J. Quirk, Prefrontal control of fear: more than just extinction. Current opinion in neurobiology, 2010. 20(2): p. 231–235.

42. Baizer, J.S., L.G. Ungerleider, and R. Desimone, Organization of visual inputs to the inferior temporal and posterior parietal cortex in macaques. Journal of Neuroscience, 1991. 11(1): p. 168–190.

43. Usrey, W.M. and H.J. Alitto, Visual functions of the thalamus. Annual review of vision science, 2015. 1(1): p. 351–371.

44. Baumann, O., et al., Consensus paper: the role of the cerebellum in perceptual processes. The Cerebellum, 2015. 14: p. 197–220.

45. Silverstein, D.N. and M. Ingvar, A multi-pathway hypothesis for human visual fear signaling. Frontiers in systems neuroscience, 2015. 9: p. 101.

46. Rahman, S.S., et al., Differential contribution of anterior and posterior midcingulate subregions to distal and proximal threat reactivity in marmosets. Cerebral Cortex, 2021. 31(10): p. 4765–4780.

47. Furl, N., et al., Top-down control of visual responses to fear by the amygdala. Journal of Neuroscience, 2013. 33(44): p. 17435–17443.

48. Maier, S., et al., Clarifying the role of the rostral dmPFC/dACC in fear/anxiety: learning, appraisal or expression? PloS one, 2012. 7(11): p. e50120.

49. Cheng, D.T., et al., Human amygdala activity during the expression of fear responses. Behavioral neuroscience, 2006. 120(6): p. 1187.

50. Dodhia, S., et al., Modulation of resting-state amygdala-frontal functional connectivity by oxytocin in generalized social anxiety disorder. Neuropsychopharmacology, 2014. 39(9): p. 2061–2069.

51. Stujenske, J.M. and E. Likhtik, Fear from the bottom up. Nature neuroscience, 2017. 20(6): p. 765–767.

52. Morawetz, C., et al., Changes in effective connectivity between dorsal and ventral prefrontal regions moderate emotion regulation. Cerebral Cortex, 2016. 26(5): p. 1923–1937.

53. Cha, J., et al., Circuit-wide structural and functional measures predict ventromedial prefrontal cortex fear generalization: implications for generalized anxiety disorder. Journal of Neuroscience, 2014. 34(11): p. 4043–4053.

54. Lee, S. and D.K. Lee, What is the proper way to apply the multiple comparison test? Korean journal of anesthesiology, 2018. 71(5): p. 353–360.

